# Conformational ensemble-based framework enables rapid development of Lassa virus vaccine candidates

**DOI:** 10.1101/2024.11.21.624760

**Authors:** Nitesh Mishra, Gabriel Avillion, Sean Callaghan, Charlotte DiBiase, Jonathan Hurtado, Nathan Liendo, Sarah Burbach, Terrence Messmer, Bryan Briney

## Abstract

Lassa virus (LASV), an arenavirus endemic to West Africa, poses a significant public health threat due to its high pathogenicity and expanding geographic risk zone. LASV glycoprotein complex (GPC) is the only known target of neutralizing antibodies, but its inherent metastability and conformational flexibility have hindered the development of GPC-based vaccines. We employed a variant of AlphaFold2 (AF2), called subsampled AF2, to generate diverse structures of LASV GPC that capture an array of potential conformational states using MSA subsampling and dropout layers. Conformational ensembles identified several metamorphic domains—areas of significant conformational flexibility—that could be targeted to stabilize the GPC in its immunogenic prefusion state. ProteinMPNN was then used to redesign GPC sequences to minimize the mobility of metamorphic domains. These redesigned sequences were further filtered using subsampled AF2, leading to the identification of promising GPC variants for further testing. A small library of redesigned GPC sequences was experimentally validated and showed significantly increased protein yields compared to controls. Antigenic profiles indicated these variants preserved essential epitopes for effective immune response, suggesting their potential for broad protective efficacy. Our results demonstrate that AI-driven approaches can predict the conformational landscape of complex pathogens. This knowledge can be used to stabilize viral proteins, such as LASV GPC, in their prefusion conformation, optimizing them for stability and expression, and offering a streamlined framework for vaccine design. Our deep learning / machine learning enabled framework contributes to global efforts to combat LASV and has broader implications for vaccine design and pandemic preparedness.

## INTRODUCTION

Proteins are inherently flexible and dynamic, with their conformational distributions influenced by temperature, solution conditions, and interactions with ligands or other proteins. Viruses with type I fusion machinery, such as HIV-1, Influenza, RSV, and LASV, have evolved conformationally plastic fusion glycoproteins known for their metamorphic nature, complicating the prediction and stabilization of their conformational states^1^. While high-resolution crystal structures can provide accurate models of dominant conformations in specific environments, these structures vary under different conditions. Artificial intelligence (AI)-based models have shown high accuracy^2,3^ but with their default parameters, are limited in their ability to account for ligands, covalent modifications, environmental factors, and protein-protein interactions, as well as multiple conformations. Recent studies have shown that AlphaFold2 (AF2) with alterations to various parameters and multiple sequence alignment (MSA) depths can predict conformational changes from sequence data alone^4–6^. These studies used subsampling and/or clustering of MSAs to adjust co-evolutionary signals across different structural domains. While the use of AF2 to sample the conformational space of proteins with few metamorphic domains has been demonstrated, a comprehensive framework for challenging targets like viral fusion proteins is lacking.

Lassa virus (LASV), a highly pathogenic arenavirus endemic to West Africa and the etiologic agent of Lassa Fever (LF), infects hundreds of thousands of people each year and presents a critical threat to global public health^7–10^. The increasing incidence and widening geographic risk zone of LASV infections highlight the urgent need for effective clinical countermeasures^11–13^. Central to vaccine and therapeutic development is the LASV glycoprotein complex (GPC), which facilitates viral entry into host cells and is the only known target of neutralizing antibodies (nAbs)^14–16^. The LASV GPC is composed of three subunits: the receptor-binding GP1, the membrane fusion GP2, and the unique stable signal peptide (SSP), which plays a critical role in GPC maturation and function. GP1 is responsible for host cell receptor binding, while GP2 facilitates membrane fusion during viral entry^17–19^. The SSP, common to all arenaviruses but distinct from other viruses utilizing class I fusion machinery, remains non-covalently associated with GP2, acting as a chaperone for proper folding and aiding in the regulation of fusion activity^20–22^. This unique structural arrangement adds to the complexity of stabilizing the GPC in its pre-fusion state for vaccine development. The inherent instability and conformational flexibility of the LASV GPC pose significant challenges to efficient recombinant expression of soluble, pre-fusion-stabilized GPC trimers, impeding vaccine development and monoclonal antibody (mAb) discovery efforts^15,23,24^. LASV nAb cocktails are protective when delivered prophylactically or therapeutically^25–27^, strongly suggesting that vaccine-induced nAbs, if elicited at sufficiently high titers, will be similarly protective. Thus, the rational design of an optimized GPC construct suitable for both immunization and nAb discovery is of paramount interest.

Previous studies have aimed to stabilize the LASV glycoprotein complex (GPC) by introducing disulfide bonds between the GP1 and GP2 subunits, resulting in constructs like GPCysR4 and GPCysRRLL^15,23^. These soluble engineered proteins maintain the GPC in its pre-fusion state, which is crucial for eliciting neutralizing antibody responses. GPCysRRLL retains the native SKI-1 cleavage site (RRLL), essential for receptor binding and recognition by neutralizing antibodies such as 8.9F^14,15^. In GPCysR4, the SKI-1 cleavage site was substituted with a furin-cleavable sequence (RRRR) to enhance processing efficiency. Further optimization by adding trimerization domains, like I53-50A^28^ and T4-fibritin^24^, has been explored to improve the stability and expression of these constructs, facilitating the formation of soluble GPCs. Although the development of a pre-fusion-stabilized GPC was an important first step for recombinant protein vaccine design, several challenges persist. Pre-fusion GPC trimers express poorly and rapidly dissociate into individual monomers^15,23,28,28^, leading to the loss of nAb epitopes and exposure of immunodominant non-nAb epitopes on the non-glycosylated GPC interior.

Given the conformational dynamism of the LASV GPC, we reasoned that an AI/ML framework capable of predicting the diverse conformational ensembles of this glycoprotein, identifying the triggering residues and/or domains, and reengineering those with residues that can stabilize the vaccine amenable pre-fusion conformation would aid in rapid vaccine development. This framework would provide critical insights into metamorphic domains—regions capable of adopting different secondary structures depending on the conformational state^5^. The knowledge gained can be used to stabilize these domains, maintain the antigenically preferred pre-fusion state, limit the exposure of non-neutralizing antibody epitopes, and design vaccine immunogens capable of reliably eliciting protective neutralizing antibody responses. This study aimed to address several critical challenges in stabilizing the LASV GPC for vaccine development and nAb discovery. Using an integrative AI-driven pipeline, including subsampled AlphaFold2^6^ and ProteinMPNN^29^, we identified crucial metamorphic domains within the LASV GPC that contribute to its conformational flexibility and targeted these regions for stabilization. We successfully engineered GPC variants optimized for expression and stability in their pre-fusion state, preserving key neutralizing antibody epitopes. These redesigned proteins exhibited significantly improved yields and retained immunogenic properties, suggesting their potential as vaccine candidates. Our approach offers a novel framework for tackling conformationally dynamic viral proteins, with promising implications for LASV vaccine design and broader applications in viral immunogen engineering.

## RESULTS

### Subsampled AF2 Predicts Conformational Ensembles of Lassa Virus Glycoprotein Complex

AF2’s default parameters restrict the heterogeneity of the predicted structures. To ensure a thorough exploration of the entire conformational landscape, we employed a sampling approach involving four MSA subsamples per pass (with each subsample providing alternate co-evolutionary information), three independent seeds, and five models per seed yielding 60 models per analysis (**FIGURE 1A**). To further enhance the diversity of predicted structures, we enabled dropout layers. Dropout layers encourage the generation of alternative solutions by randomly deactivating certain weights during training. By extending this principle to inference, we augmented the diversity of predicted structures, thus enabling a systematic sampling of the conformational space. This comprehensive sampling approach facilitated robust analyses of LASV GPC conformational diversity.

**Figure 1.**
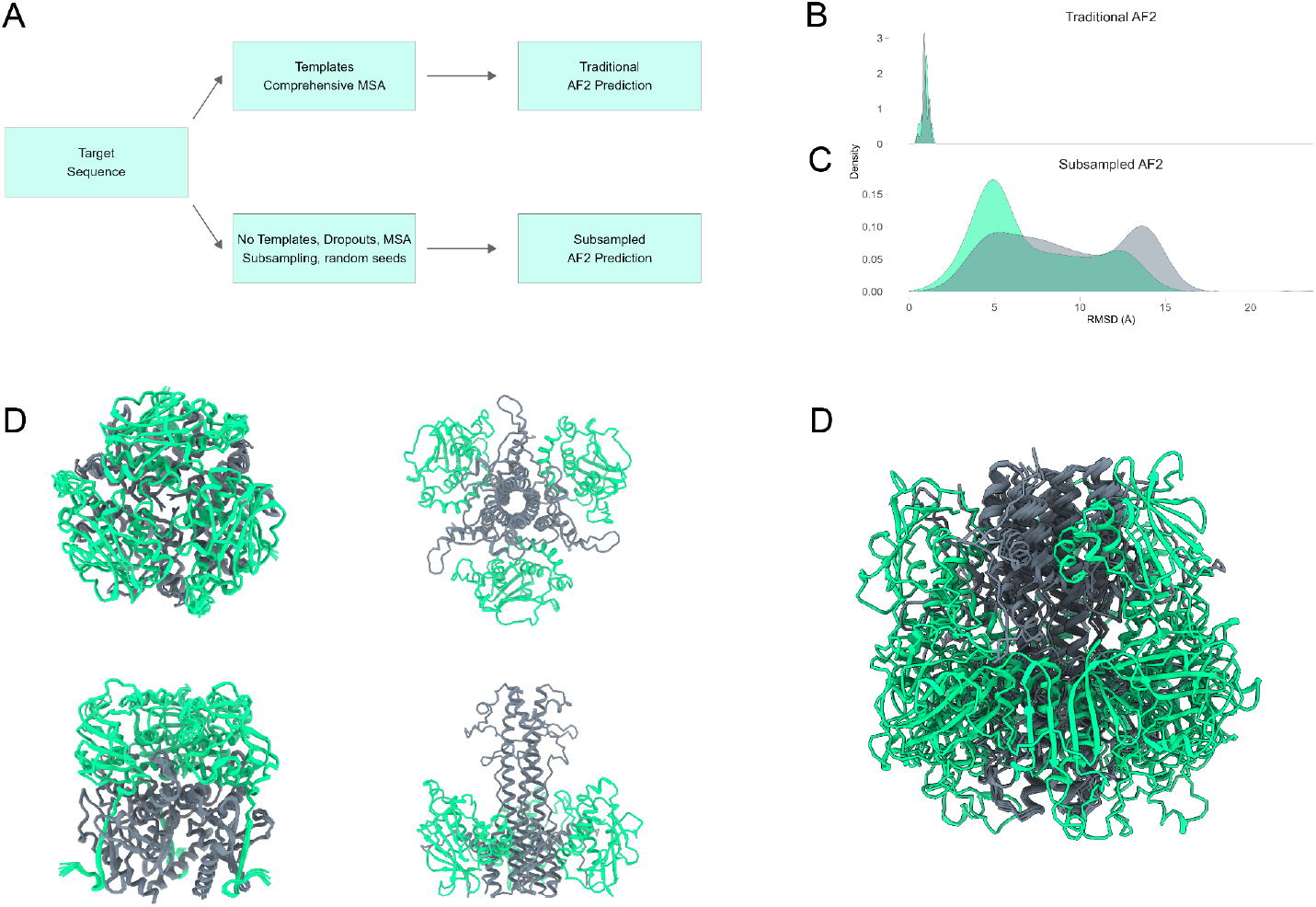
Sampling of LASV GPC Conformational Diversity Using Subsampled AF2. (A) Workflow illustrating the sampling strategy employed to enhance conformational heterogeneity. (B) Structural predictions with default AF2 parameters predominantly yield the pre-fusion conformation, with limited exploration of alternative conformations. (C) Subsampled AF2 pipeline captures a broader spectrum of LASV GPC conformations. (D-F) The AF2 predicted ensemble spans pre-fusion conformation while subsampled AF2 ensembles span varying conformations of post-fusion GPC.

When using AF2 with its default settings, which involve a deep MSA and template bias, the pre-fusion conformation of LASV GPC dominates the prediction set. By relying heavily on structural templates, AF2’s predictions tend to converge towards a single conformation, disregarding the diverse range of conformational forms that LASV GPC may adopt (**FIGURE 1B**). As anticipated, with the subsampled AF2 pipeline, a broader spectrum of LASV GPC conformations was observed (**FIGURE 1C**). Models generated without templates generally exhibited low predicted Local Distance Difference Test (pLDDT) scores, except for the receptor-binding domain, which showed high pLDDT values at specific residue positions (e.g., Josiah residue numbers) (**SUPPLEMENTARY**). pLDDT values correlate inversely with protein structural flexibility; metamorphic viral fusion proteins generally exhibit lower pLDDT scores, while proteins with well-defined secondary structures, particularly those rich in helices and strands, tend to produce higher pLDDT scores. The predicted ensemble spans known conformational states of LASV (pre- and post-fusion as well as cleaved versus uncleaved) without generating unphysical or severely misfolded structures (**FIGURE 1D-F**). This remarkable coverage of the conformational space underscores the efficacy of AF2 in sampling intermediate states of viral type I fusion machinery, offering valuable insights into domains governing LASV GPC conformational dynamics.

### Mapping LASV Metamorphic Domain Dynamics

The conformational structure ensemble generated by AF2 through MSA subsampling was next used to investigate the structural dynamics of the LASV GPC in its uncleaved and cleaved stages. Our analysis focused on regions exhibiting significant structural deviations across ensemble models, which we label as metamorphic domains and are characterized by marked conformational flexibility. Core domains within both GP1 and GP2 were consistently modeled within 4-5 recycles, beyond which the models displayed diverse loop conformations (**SUPPLEMENTARY**). This variability underscores the intrinsic conformational plasticity inherent in the LASV GPC domains. For uncleaved LASV GPC, the majority of the conformational plasticity was seen within the N-terminal fusion peptide (NFP), internal fusion loop (IFL) and heptad repeat 1 (HR1) domains (**FIGURE 2A**). In contrast, for the cleaved GPC, the majority of the HR1 was in its six-helix bundle (6HB) form and had the least conformational plasticity. In cleaved GPC, most of the conformational plasticity was seen in the α5 helix of GP1, NFP, IFL and T-loop region of GP2 (**FIGURE 2B**).

**Figure 2.**
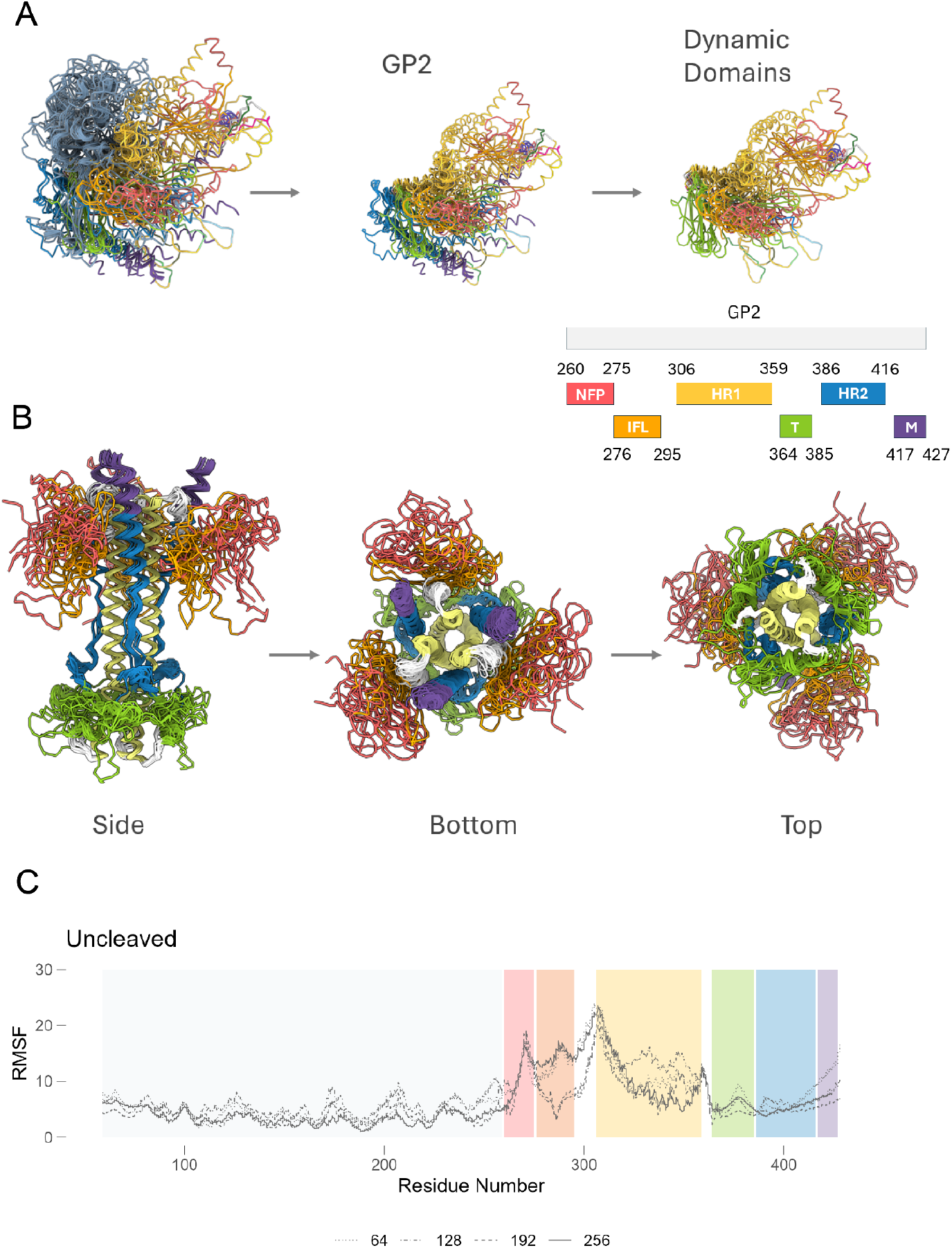
Conformational ensembles predict metamorphic domains within Lassa GPC. (A) Superimposed predicted structures for uncleaved monomers for Lassa josiah strain highlight conformational plasticity within NFP, IFL and HR1 domains. (B) Superimposed predicted structures of post-fusion conformation highlighting conformational plasticity within NFP, IFL and T-loop during the post-fusion stage. (C) Trajectory analysis for all predicted structures showing conformationally dynamic regions of lassa GPC at different MSA subsamples (as denoted by varying lines). The known domains of GP2 are color-coded: NFP (red); IFL (orange); HR1 (yellow); T-loop (green); HR2 (blue); MPER (purple).

To gain deeper insight into the dynamic behavior of LASV GPC, we conducted trajectory analysis on a subset of AF2 predicted models. Specifically, trajectories were constructed from the final models generated using three seeds and four MSA subsamples to examine conformational transitions and identify regions prone to instability. The transition of GP1 from pre-to post-fusion involves moderate conformational changes, whereas GP2’s transition entails significant backbone rearrangements. In GP1, apart from disordered loops, the α5 helix exhibited the highest conformational plasticity. In GP2, the HR1 and HR2 regions formed typical six-helix bundles, with the T-loop linking both domains sampling an extensive conformational landscape. The most pronounced conformational plasticity was observed in the NFL and IFL of the fusion domain, indicating the protein’s capacity for structural adaptation and refolding under varying environmental conditions. These regions, characterized by high Root Mean Square Fluctuation (RMSF) values, are potential targets for stabilization efforts aimed at enhancing the overall structural integrity of the protein complex.

### ProteinMPNN-Driven LASV GPC Optimization

Stabilization of the pre-fusion conformation of viral surface glycoproteins is primarily achieved by inhibiting the release of the fusion peptide or by disrupting the formation of the coiled-coil structure typical of the post-fusion state^30–32^. Two widely used and effective strategies include designing disulfide bonds at regions that undergo significant refolding and introducing proline substitutions to hinder the formation of the central post-fusion helices. Using the subsampled AF2 conformational ensemble, with a particular focus on residues exhibiting high conformational dynamics within the metamorphic domains, we employed ProteinMPNN, a structure-to-sequence predictor^29^, to strategically redesign these pivotal residues. Based on the conformational ensemble data, key regions for stabilizing the pre-fusion conformation include the GP1-GP2 interface (α5 helix, which interfaces with GP2 between the α8 and α9 helices) (**FIGURE 3A**). Additionally, several residues that formed stabilizing interactions in the post-fusion 6HB conformational ensemble were also considered: K320, R325, K327, and E329 (**FIGURE 3B**). These targeted interventions were anticipated to reinforce the structural stability of LASV GPC in its perfusion conformation, contributing to the development of more robust immunogens.

**Figure 3.**
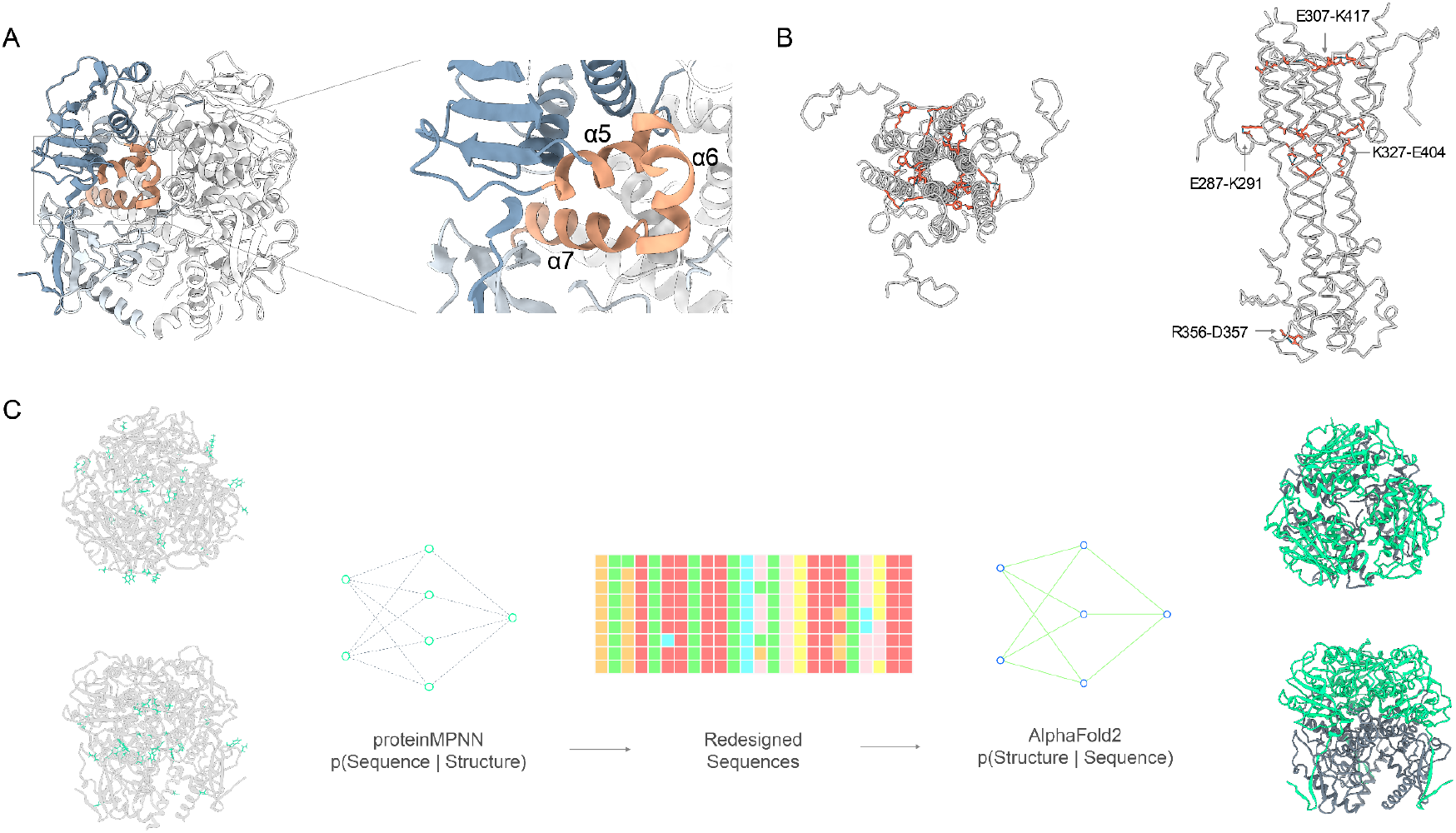
Stabilization Strategies for Pre-Fusion LASV Glycoprotein Conformation. (A) GP1-GP2 interface highlighted for stabilization, showing the α5 helix (from GP1) interaction with α6 and α7 helices (from GP2 subunit). (B) Residues (K320, R325, K327, E329) from the post-fusion 6HB conformational ensemble that form stabilizing interactions, illustrating their importance in pre-fusion stability. (C) Overview of the ProteinMPNN design protocol applied to the pre-fusion LASV GPC structure (PDB: 7PVD), showcasing the redesign strategy to enhance structural stability while preserving mAb binding epitopes and N-linked glycosylation sites. The generation of 2,306 sequences from varying temperature parameters and models is illustrated, leading to a refinement process that identified the top 24 designs for experimental evaluation.

By utilizing these conformationally dynamic and post-fusion stabilizing residues as seeds for ProteinMPNN driven redesign, we aimed to introduce strategic modifications that would mitigate the observed structural variability and enhance protein stability. We applied the ProteinMPNN design protocol (**FIGURE 3C**) to the pre-fusion LASV GPC structure (PDB: 7PVD), after filling missing loops within the crystal structure using AF2. ProteinMPNN generated a diverse array of sequences, each representing a variant of LASV GPC with potentially improved stability and expression. Additional constraints were imposed during sequence design to ensure the preservation of key mAb binding epitopes and all 11 potential N-linked glycosylation sites as well as minimizing backbone redesign to maintain structural integrity. A total of 2,306 sequences were generated using ProteinMPNN, incorporating variations across three different temperature parameters and three separate models. To refine the design space, combinatorial sequence design was performed using consensus residues identified across multiple temperatures and models. The top 200 sequences were further filtered using AF2 based on pLDDT scores and Cα root-mean-square deviation (RMSD) to the pre-fusion LASV structure (PDB: 7PVD). We selected the top 24 designs for experimental characterization. Additionally, we included 24 low-scoring designs with unfolded or disordered fusion domains and HR1 regions as negative controls.

### Antigenic validation of redesigned LASV GPC sequences

The 48 redesigned sequences (24 high pLDDT designs, 24 low pLDDT designs) were assessed for generating well-ordered, soluble LASV envelope trimer mimetics. Given the inherent dissociation tendency of GP1 and GP2, we incorporated a flexible linker between these two domains to maintain structural integrity by covalently linking the metastable subunits to counteract spontaneous dissociation. The linker was designed to mimic the natural flexibility found in cleaved viral glycoproteins, thereby preserving the native-like conformational dynamics essential for functional integrity. The linker length was selected based on the Cα distance between residues L259 and G260 in the cleaved pre-fusion structure (PDB: 7PUY), corresponding to approximately 6.5 residues in a fully extended peptide backbone, as informed by previous literature^33,34^. Notably, our strategy diverges from conventional LASV GPC engineering methods by omitting trimerization domains or scaffolds at the C-terminus^15,24,28^ and intra- and inter-disulfide linkage of the two subunits^23,24^.

All 24 designs with high pLDDT AF2 predictions yielded significantly higher total soluble protein (up to a 4.1-fold increase) compared to GPCysR4-I53A (**FIGURE 4A-B**). Conversely, none of the 24 control designs with low pLDDT or disordered domains expressed well. The CysR4 and CysRRLL constructs did not express at significant levels in small-scale transfections, which aligns with their previously determined expression level in large-scale settings^24,28^. This outcome underscores the critical role of protein stability and structural integrity in efficient protein expression. Importantly, the absence of appended trimerization domains or scaffolds at the C-terminus did not hinder the expression levels of the redesigned LASV GPC variants, highlighting the effectiveness of our redesign strategy in enhancing protein stability and, by extension, expression yield.

**Figure 4.**
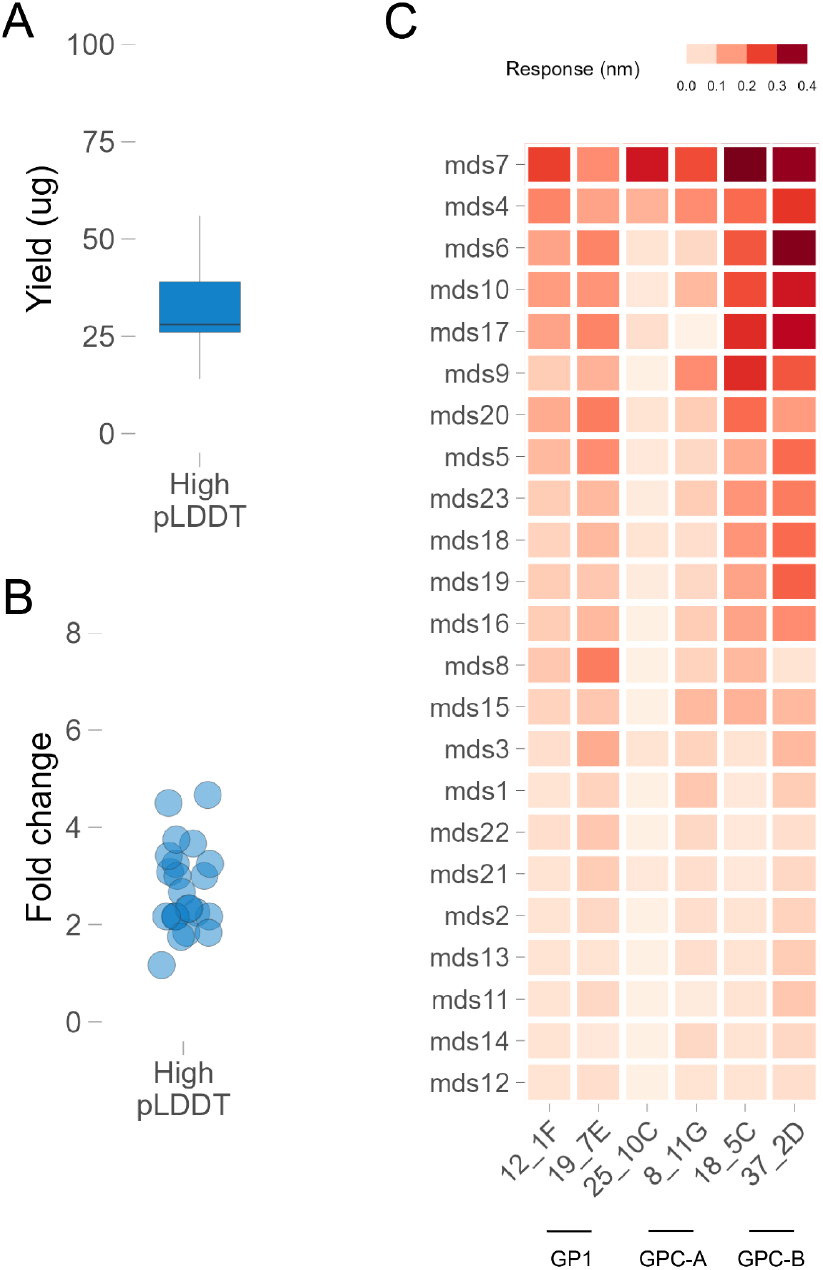
Antigenic characterization of ML designed LASV GPC constructs. (A) Protein yield of 24 lead designs (with high pLDDT confidence predictions). (B) Fold change compared to GPC-CysR4-I53A for designs from panel A. (C) Antigenic profile of all 24 designs with a panel of conformation specific monoclonal antibodies.

To verify that the redesigned LASV GPC variants’ enhanced expression did not compromise functionality, we conducted comprehensive antigenic characterization. Initial functional screening involved evaluating antigenicity using a panel of antibodies specific to various conformational epitopes of pre-fusion LASV GPC. We utilized seven previously described nAbs representing four distinct epitope classes: 12.1F and 19.7E targeting GP1-A, 25.10C and 8.11G targeting GPC-A, 37.2D and 18.5C targeting GPC-B, and 8.9F targeting GPC-C^14^. The GP1-A group binds to GP1 even in the absence of GP2, while the other groups recognize only the fully assembled GPC (containing both GP1 and GP2). Most antibodies, with the exception of the GPC-C antibody 8.9F, showed measurable binding (**FIGURE 4C**), indicating that the stabilized trimer retained antigenic properties similar to the GPCysR4-I53-50A construct, despite the absence of additional trimerization domains or scaffolds. This binding profile suggests that the flexibly linked redesigned trimer retains most of the epitopic landscapes of a pre-fusion conformation crucial for effective immune recognition, which is essential for both therapeutic and vaccine applications. Furthermore, the preserved binding to multiple nAbs indicates the potential for broad protective efficacy, as these antibodies target diverse and critical epitopes on the GPC.

## DISCUSSION

In this study we leveraged advanced deep learning techniques, specifically AlphaFold2^3^ and ProteinMPNN^29^, and developed a conformational ensemble-based framework to stabilize and enhance the expression and antigenicity of the Lassa virus GPC^7,9,10,23^, a critical target for vaccine development. This *in-silico* approach enabled us to sample an extremely large mutational space and rapidly prioritize a small library of redesigned GPC sequences, ultimately identifying variants that exhibited improved structural stability and solubility while maintaining key immunogenic epitopes. The application of AF2 for generating conformational ensembles^5,6^ allowed us to identify metamorphic domains within the LASV GPC, which are regions with significant conformational plasticity, highlighting that AF2, augmented by targeted sampling strategies, can effectively predict conformational ensembles of complex viral proteins like LASV GPC. This capability is critical for understanding the structural basis of viral fusion^17–19^ and for guiding the design of more effective vaccines and therapeutics targeting the pre-fusion state of viral glycoproteins. Stabilizing these domains in the prefusion state is crucial for vaccine discovery, preserving neutralizing antibody epitopes and preventing the exposure of immunodominant non-neutralizing epitopes. This targeted stabilization is a significant advancement, as it enhances the potential immunogenicity of the GPC, making it a more effective candidate for vaccine development. The increased levels of soluble protein yield observed in our redesigned GPC constructs, coupled with their retention of crucial epitopic landscapes, underscore the potential of these variants for generating a robust immune response. This finding is particularly important as it suggests that our approach can lead to the development of vaccines with broad protective efficacy against LASV.

One particular advantage of our approach is the absence of any disulfide linkage or structural trimerization / foldon scaffolds that can potentially generate spurious immune responses^24,28^. This approach led to the formation of soluble trimeric GPC that binds all, except one, tested neutralizing antibodies without the need for additional trimerization domains or conformation-locking antibodies. This design strategy demonstrates potential in enhancing the stability and expression of LASV envelope proteins, paving the way for advancements in vaccine development and therapeutic antibody design. Furthermore, the ability to maintain functional antigenicity without structural scaffolding offers a streamlined pathway to produce vaccine candidates, potentially reducing manufacturing complexities and costs. This strategy is readily generalizable to other viral glycoproteins^30–32,34^, highlighting its broad applicability in virology and immunogen design.

Despite the promising results, there are several limitations to our study. First, while our computational models predicted increased stability and solubility of the redesigned GPC variants, these findings need to be validated through extensive in vivo experiments to confirm their immunogenicity and protective efficacy. The complexity of the immune system and the potential for unforeseen interactions mean that computational predictions, while valuable, are not a substitute for empirical testing. Second, our study focused on the LASV GPC, which, although critical, represents just one component of the virus. The efficacy of a potential vaccine will also depend on other viral and host immune factors, such as the induction of robust T cell responses, which were beyond the scope of this study. Additionally, while the principles remain the same, the generalizability of our framework to other viral proteins and pathogens remains to be fully explored. Third, while our approach successfully identified GPC variants with enhanced expression and stability, the long-term stability and efficacy of these variants under physiological conditions require further investigation as the dynamics of protein folding, and potential post-translational modifications in vivo could impact the stability and immunogenicity of the redesigned GPC. Future research should aim to validate these computationally stabilized trimers in animal models to assess the immunogenicity and protective efficacy of the redesigned GPC variants. Such studies will be crucial in determining the potential of these variants as vaccine candidates. Additionally, exploring the application of our framework to other viral proteins and pathogens will provide broader insights and contribute to the development of vaccines and therapeutics for a wide range of diseases. Further refinement of the conformational ensemble-based framework could involve integrating additional computational tools and experimental data to enhance the information gained from ensemble predictions. Expanding our understanding of the structural dynamics of viral proteins will be essential for improving the design of immunogens and therapeutic antibodies.

In conclusion, our study demonstrates that an AI-driven framework can effectively predict and stabilize the conformational states of complex viral proteins like LASV GPC. The insights gained from this research contribute to the broader effort of developing effective vaccines and therapeutic interventions for LASV, addressing a pressing global health challenge. Continued advancements in computational modeling and experimental validation hold promise for accelerating the discovery and development of vaccines and treatments for emerging infectious diseases.

## METHODS

### Conformational Ensemble Generation

We used a custom implementation of colabfold^35,36^ 1.5.3 to predict the cleaved and uncleaved structures of LASV GPC using the mmseqs^37^ server for MSA generation. MSAs were subsampled at following max_seq:extra_seq ratios (256:512, 192:384, 128:256, 64:128).

### MPNN parameters

For redesigning residues stabilizing LASV GPC in its perfusion confirmation, we used a custom implementation of proteinMPNN, a structure-to-sequence ML model with model version V_48_020. 512 redesigned sequences were generated at sampling temperature of 0.001, 0.01, 0.1 each with the homooligomer constraint using an AF2 model of LASV GPC as the structure backbone. The backbone was fixed while the following positions within the GPC were used for redesigning (**POSITIONS 61, 82, 135, 248 and 254** within GP1, **262, 264, 283, 287, 312, 322, 327, 329, 336, 337, 338, 341, 345, 347 and 350** within GP2).

### Protein Expression and purification

Initial characterization of all variants was performed with small 50 ml 293F cultures. 293F cells in 100 ml erlenmeyer flasks were transfected with 40 ug of redesigned LASV GPC constructs and control GPC R4 and RRLL constructs. FIve days post-transfection, supernatants were harvested and protein was purified from the supernatant using GNL affinity columns. The purified proteins were buffer exchanged with PBS and concentrated down using Amicon 30K filters.

### Antigenic Screening

BLI was performed with12.1F and 19.7E targeting GP1-A, 25.10C and 8.11G targeting GPC-A, 37.2D and 18.5C targeting GPC-B, and 8.9F targeting GPC-C. IgGs were immobilized on ProA sensors (Sartorius) to a signal of 1.0 nM using an Octet Red96 instrument (ForteBio). The immobilized IgGs were then dipped in the running buffer (PBS, 0.1% BSA, 0.02% Tween20, pH 7.4) followed by redesigned 500 nM of LASV GPC trimer, or running buffer. Following a 120 s association period, the tips were dipped into the running buffer and dissociation was measured for 240 s.

### Human LASV antibodies expression

IgG heavy and light chain sequences for lassa GPC targeting antibodies from Robinson et.al., were constructed and cloned in AbVec Vector. Paired heavy and light chain plasmids were co-transfected in Expi293 cells in a !:1 ratio using Fectopro transfection reagent. Monoclonal IgGs were purified from the culture supernatant five days post-transfection with Protein-A sepharose per manufacturer’s instructions.

### Structure Visualization

For visualizing aligned structures, generating superimposed ensemble, color structures by AlphaFold2 confidence score (pLDDT), UCSF ChimeraX^38^ (version 1.8) was used.

### Code

Raw data used in this study will be made available on Zenodo and all the code used in this study is available on GitHub (https://github.com/brineylab/lasv-gpc-stabilization).

## AUTHOR CONTRIBUTIONS

Conceptualization: NM, BB

Code and modeling: NM, GA, SC, SB

Protein expression and testing: NM, GA, SC, CD, TM Reagents: JH, NL

Data analysis: NM, SC

Manuscript preparation and revisions: NM, GA, SC, CD, JH, NL, SB, TM, BB

